# Cryptic Promoter Activation Drives *POU5F1* (*OCT4*) Expression in Renal Cell Carcinoma

**DOI:** 10.1101/379198

**Authors:** Kyle T. Siebenthall, Chris P. Miller, Jeff D. Vierstra, Julie Mathieu, Maria Tretiakova, Alex Reynolds, Richard Sandstrom, Eric Rynes, Shane J. Neph, Eric Haugen, Audra Johnson, Jemma Nelson, Daniel Bates, Morgan Diegel, Douglass Dunn, Mark Frerker, Michael Buckley, Rajinder Kaul, Ying Zheng, Jonathan Himmelfarb, Hannele Ruohola-Baker, Shreeram Akilesh

## Abstract

Transcriptional dysregulation drives cancer formation but the underlying mechanisms are still poorly understood. As a model system, we used renal cell carcinoma (RCC), the most common malignant kidney tumor which canonically activates the hypoxia-inducible transcription factor (HIF) pathway. We performed genome-wide chromatin accessibility and transcriptome profiling on paired tumor/normal samples and found that numerous transcription factors with a RCC-selective expression pattern also demonstrated evidence of HIF binding in the vicinity of their gene body. Some of these transcription factors influenced the tumor’s regulatory landscape, notably the stem cell transcription factor *POU5F1* (*OCT4*). Unexpectedly, we discovered a HIF-pathway-responsive cryptic promoter embedded within a human-specific retroviral repeat element that drives *POU5F1* expression in RCC via a novel transcript. Elevated *POU5F1* expression levels were correlated with advanced tumor stage and poorer overall survival in RCC patients. Thus, integrated transcriptomic and epigenomic analysis of even a small number of primary patient samples revealed remarkably convergent shared regulatory landscapes and a novel mechanism for dysregulated expression of *POU5F1* in RCC.

## Introduction

Development of new therapeutic strategies for cancer treatment depends on identification of critical mechanisms and pathways utilized by tumor cells. Numerous insights have been gleaned from large tumor consortium programs such as The Cancer Genome Atlas (TCGA), which has extensively catalogued somatic mutations and selected phenotypic features from thousands of tumor and normal tissue samples across a variety of human cancers. To some extent, insights from such broad-based studies are intrinsically limited by tumor heterogeneity (including presence of non-tumor cell types) and general sample variability, which may collectively obscure sensitive and robust detection of subtle changes in cellular pathways such as transcription factor regulatory networks that define and govern the malignant state (Stergachis et al. 2013). Epigenomic mapping of tumors in large consortium-driven projects has generally focused on DNA methylation analysis (TCGA, Roadmap Epigenomics Project) and targeted histone modification profiling using ChIP-seq (Roadmap). These systematic approaches leverage the fact that patterns of regulatory DNA (e.g. promoters, enhancers, insulators) activation and organization are extensively disrupted in cancer (Stergachis et al. 2013; Polak et al. 2015). Generic identification of regulatory DNA is best achieved by open chromatin profiling methods such as DNase-seq (Boyle et al. 2008) and ATAC-seq (Buenrostro et al. 2013). However, the complexity of these deep epigenomic mapping methods has focused their initial application to mouse tissues (Yue et al. 2014), cultured human cell lines (Thurman et al. 2012), whole adult and fetal human tissues (Kundaje et al. 2015), hematopoietic neoplasms (where both malignant and normal cells of origin are readily obtained (Corces et al. 2016; Qu et al. 2017)), and a limited number of epithelial malignances (Polak et al. 2015). When deploying sensitive epigenomic methods, matched normal tissues of origin provide the best control for patient genotype and environmental exposure but are often discarded or unavailable at the time of tumor resection. Taken together, these hurdles have limited the characterization of primary human epithelial malignancies together with their patient-matched normal cells-of-origin.

In this regard, clear cell renal cell carcinoma (RCC), the most common and lethal kidney malignancy, is an ideal model cancer system for high-resolution functional genomic analyses for several reasons. First, RCC tissues are readily available since the standard of care is surgical removal of the often-large tumor mass, frequently with plentiful adjacent, non-neoplastic tissue. Second, the tumor cells and their cells-of-origin – proximal tubule epithelial cells (Chen et al. 2016) – are readily isolated at high purity and grow well in short-term primary cultures (Cifola et al. 2011), which removes the obstacle of contaminating non-relevant cell populations. Third, the majority of spontaneously arising tumors utilize a common oncogenic pathway: stereotypic loss of chromosome 3p, resulting in loss of heterozygosity for the *VHL* tumor suppressor gene combined with inactivation of the remaining allele of *VHL* (Seizinger et al. 1988). While it is well understood that loss of functional VHL protein leads to constitutive stabilization of two DNA-binding transcription factors, hypoxia-inducible factors 1α and 2α (HIF1α, HIF2α) (Maxwell et al. 1999), the precise nature of genomic dysregulation downstream of HIF pathway activation that drives oncogenesis remains poorly understood. Given that RCC has an annual incidence of >60,000 and mortality of >14,000 in the United States alone (NCI SEER database), additional insights are urgently needed to develop new treatments.

Here, using a combination of DNase I-hypersensitivity mapping (DNase-seq) and transcriptome profiling (RNA-seq) of primary tumor and normal cell cultures derived from three patients, we uncover a high degree of concordance in the epigenomic landscape of RCC. Analyses of these high-resolution reference maps in conjunction with publicly available datasets (Cancer Genome Atlas Research 2013; Salama et al. 2015; Ricketts et al. 2018) reveal unexpected insights into the transcription factors driving genome dysregulation in RCC, notably the stem cell factor *POU5F1* (*OCT4*). This approach provides a general framework for the analysis of other solid tumors for which matched malignant and normal cells can be isolated at high purity, and greatly amplifies the utility of cancer -omics catalogs.

## Results

### RCC regulatory landscapes are highly concordant across individual tumors

Using RCC as a model system, we first sought to reduce or eliminate the contribution of non-relevant cell types by generating primary cultures of RCC and proximal tubules (cell of origin for RCC) from three patients. In culture, tumor cells were large, grew slowly and frequently contained intracellular vacuoles, typical of adenocarcinoma. In contrast, proximal tubule cells were epithelioid in morphology and grew rapidly (**Figure 1A**). Previous work has demonstrated that primary RCC cultures preserve the cytogenetic profile of their originating tumor (Cifola et al. 2011). In line with this, we found that the primary tumor cultures revealed characteristic karyotype abnormalities associated with RCC: all three patients’ tumors carried a loss of the short arm of chromosome 3 (chr3p-) and a gain of the long arm of chromosome 5 (chr5q+) (**Figure 1B and Supplemental figure 1A**). The *VHL* gene is located on chr3p, and Sanger sequencing of the remaining allele identified inactivating missense mutations in all three tumor samples (**Supplemental figure 1B**). Taken together with the loss of heterozygosity on chromosome 3p, this indicated that all three patients’ tumors were *VHL*-null, typical of the majority of sporadic RCC (Cancer Genome Atlas Research 2013).

Next, we generated high-quality DNase-seq datasets in duplicate from each patient’s primary RCC and tubule cultures. Windowed aggregation of DNase-seq tags again corroborated chromosome arm-level gains and losses delineated by conventional karyotyping (**Supplemental figure 1C**). Globally, accessible chromatin regions appear as DNase-hypersensitive sites (DHSs, called at FDR 1%) and most of these were located >5 kilobases (kb) from known transcription start sites, a feature that is typical of enhancer elements (**Supplemental figure 1D**). In parallel, we generated gene expression profiles (RNA-seq) from these cultures and compared them to TCGA RNA-seq data generated from 72 normal kidney tissues and 534 RCC specimens (Cancer Genome Atlas Research 2013). Lastly, we cross-referenced our DNase-seq and RNA-seq datasets with publicly available ChIP-seq data for HIF components (HIF1 α, HIF2α, HIF1β) from the *VHL*-null 786-O RCC cell line (Salama et al. 2015). As an example of such comparison, *STC2*, a well-known HIF-induced target gene (Law and Wong 2010), had several differentially accessible DHSs near its promoter in the RCC samples which correlated with increased *STC2* gene expression in our own data and in the larger TCGA data set (**Figure 1C**). Some of the induced DHSs near the *STC2* promoter overlapped HIF ChIP-seq peaks, consistent with HIF binding at these regulatory elements. However, other induced DHSs do not appear to be bound by HIF, implicating a role for transcription factors (TFs) in opening nuclear chromatin at these sites.

**Figure 1.**
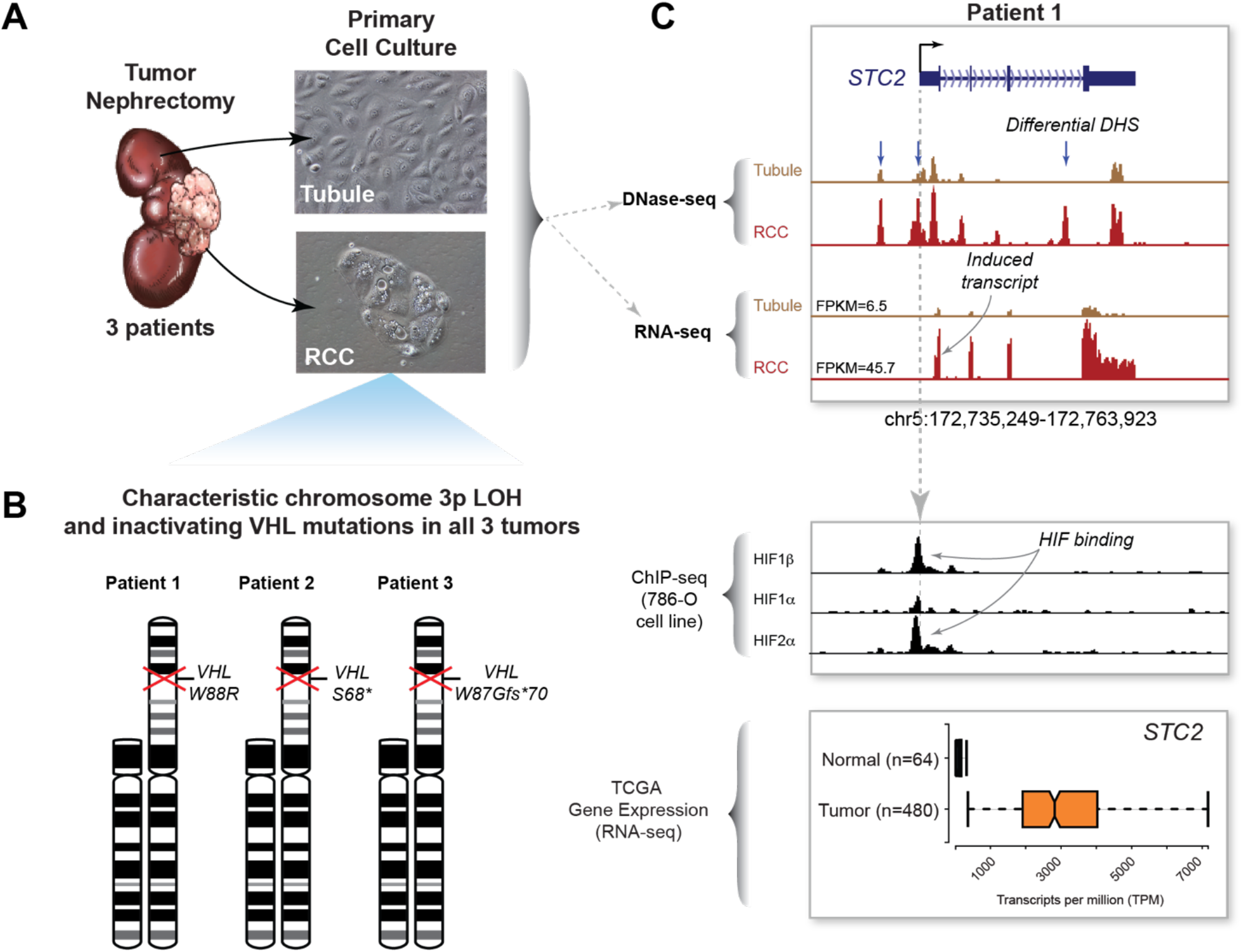
Overview of patient samples and data sets used for integrated analyses. (A) *Primary culture of tumor and matched-normal tubule cells from three patients*. Renal cell carcinoma tumor nephrectomies from three patients were used to derive primary cultures of proximal tubules and renal cell carcinoma. (B) *Cytogenetic analysis of primary tumor cultures*. Karyotype analysis of the carcinoma cultures revealed loss of the short of arm of chromosome 3 in all three patient samples. Sanger sequencing of the *VHL* gene in these same samples identified inactivating mutations in the remaining copy. (C) *Example of integrated analysis at the STC2 gene locus*. DNase-seq and RNA-seq datasets were also generated from the primary tubule and carcinoma cultures and compared to HIF ChIP-seq datasets from the 786-O renal cell carcinoma cell line and RNA-seq expression data from TCGA. *STC2*, a canonical HIF target gene, exhibits several differential DHS (blue arrows), some of which coincide with HIF binding determined by ChIP-seq. Compared to normal tubules, the *STC2* transcript is strongly induced in the primary tumor cultures and in the TCGA tumor samples.

**Supplemental Figure 1.**
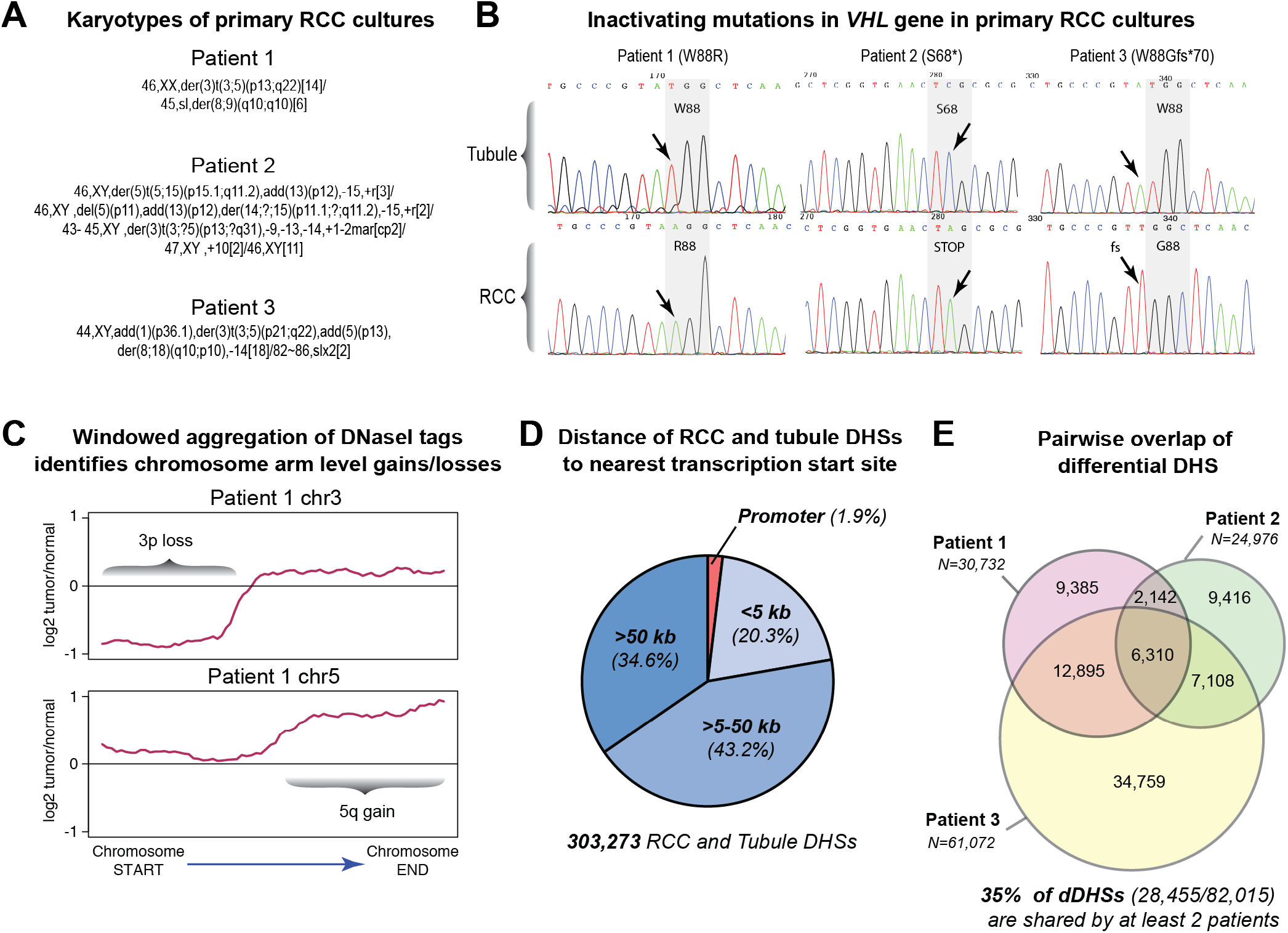
Characterization of primary RCC cultures and overview of DHS landscape. (A) *Karyotypes of primary RCC cultures*. Primary RCC cultures were submitted for G-band karyotyping at the University of Washington Cytogenetics Laboratory. Inferred karyotypes from 20 metaphase spreads are shown. All three patient tumors show characteristic loss of chromosome 3p and gain of chromosome 5q. (B) VHL *mutation status in primary tubule and RCC cultures*. Sanger sequencing of the coding regions identifies an inactivating mutation in the single copy of the *VHL* gene in all three primary RCC cultures. (C) *Chromosome arm level gains and losses are identified by DNase-seq tags*. Windowed aggregation (5Mb windows) of tags from DNase-seq datasets from the primary tubule and RCC cultures reveals arm level gains and losses, including the canonical loss of chromosome 3p and gain of chromosome 5q. (D) *DNase I-hypersensitive sites identify predominantly distal regulatory elements*. A minority of the master list DHS derived from tubule and RCC datasets localize to promoter elements (1.9%) or lie within 5 kb of a known transcription start site (20.3%). The majority (77.8%) lie >5 kb from known transcription start sites, characteristic of distal regulatory elements such as enhancers. (E) *Overlap of differential DHS identifies the shared regulatory landscape of RCC*. DHSs with differential accessibility in tumors vs. their matched tubule controls define the differentially accessible regulatory landscape of RCC. Pair wise comparisons of these differential DHS across patients reveals that ~35% of all differential DHS are shared among at least two patient samples.

Genome-wide chromatin accessibility patterns define the regulatory landscape of each primary patient sample. Globally, the regulatory landscapes of the primary tubule cultures show substantial overlap among the three profiled patients (**Figure 2A**). In contrast, while each tumor specimen retains a proportion of DHSs from its tubule of origin, the remainder of its landscape is composed of *de novo* DHSs. A proportion of these *de novo* DHSs is shared among the tumor samples, and together with the tubule-derived DHSs retained in the tumors, they define the shared regulatory landscape of RCC. The similarity of the tubule regulatory landscapes is also evident in the tight clustering of these samples in principal component analysis whereas the RCC samples (and the 786-O RCC cell line) localize to distinct positions in the regulatory space (**Figure 2B**).

After obtaining a global picture of regulatory landscape similarities based on presence or absence of individual DHS peak calls, we identified accessibility changes between each patient’s normal and tumor cells at a common set of DHSs, and then compared the behavior of those differentially accessible sites across all three patients. This analysis identified between 24,976-61,072 differential DHSs (dDHSs, FDR 1%; see Methods) in each patient (roughly split between sites with increased and decreased accessibility in tumor cells), representing ~8-20% of all sites examined (**Supplemental figure 1E**). At least 35% of these dDHSs were shared by at least 2 patients. Most strikingly, we found that 93.6-98.5% of dDHSs shared between any two patients displayed highly concordant directional accessibility changes in the tumor samples (**Figure 2C**). In total, we identified 6,080 dDHSs with concordant accessibility changes across all three patients.

The above results show that primary cultures of proximal tubules and RCC can be generated at high purity and provide an ideal platform for functional genomic methodologies. While the regulatory landscape of each patient’s tumor cells is in part unique, the shared DHSs show highly convergent accessibility changes across all three patients and therefore define the core regulatory program of RCC.

**Figure 2.**
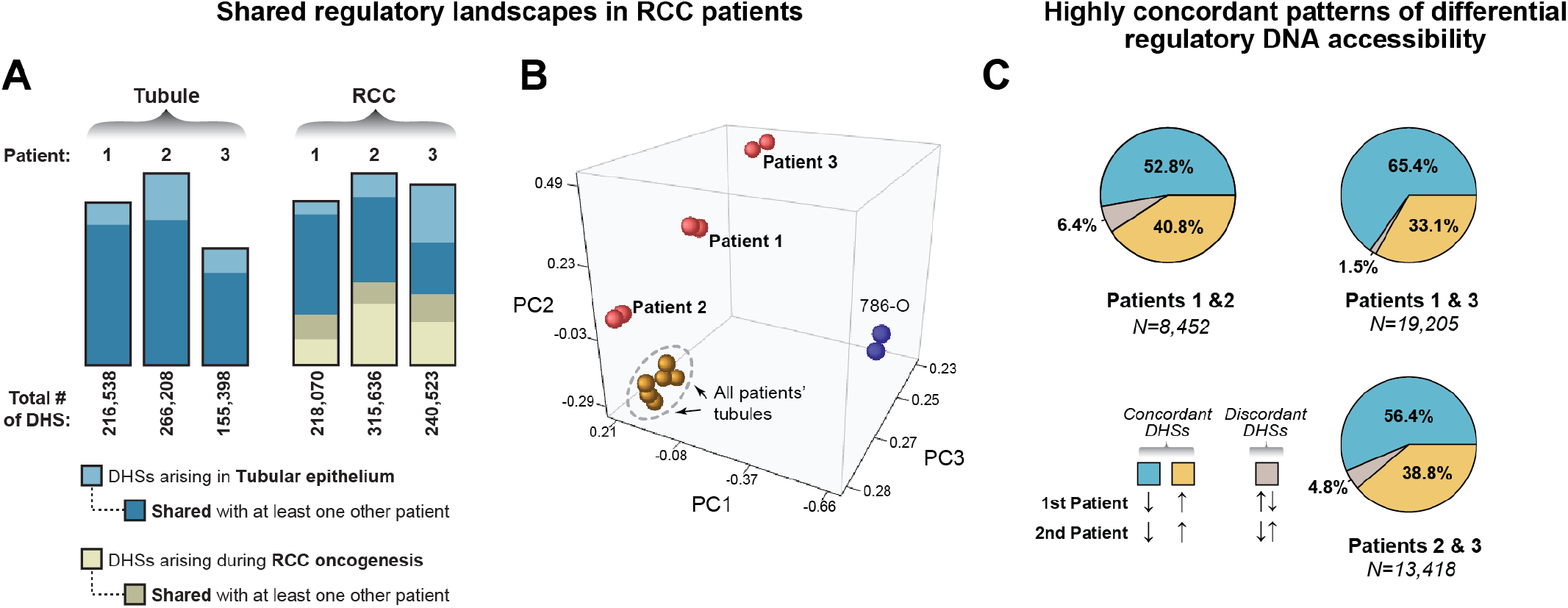
Shared regulatory landscapes in tubules and matched renal cell carcinomas from three patients. (A) *Comparison of the shared regulatory landscape among patient samples*. The three tubule samples share a significant proportion of DHSs. Each tumor’s landscape of DHSs incorporates a different fraction of DHSs from its tubule of origin and activates de novo DHSs. In the tumors, most of the tubule-derived DHSs are shared with tubule-derived DHSs from other patients. In contrast, a smaller fraction of RCC-derived de novo DHSs is shared among patient tumors. (B) *Comparison of DNase-seq data by principal component analysis*. While the tubule cultures from all three patients (brown spheres, in replicate) are tightly clustered, each tumor (red spheres, in replicate) and the 786-O cell line (blue spheres, in replicate) occupy a unique position in regulatory space. (C) *Differential DHSs show highly concordant patterns of accessibility across patient samples*. In pairwise comparisons, the shared differential DHSs are classified as concordantly upregulated in the tumor samples (gold), downregulated in the tumor samples (blue) or discordant in the two patient samples being compared (grey). The majority (>95%) of shared differential DHS show concordant up- or downregulation.

### Convergent gene expression landscapes

Examination of gene expression profiles for genes changing by >1.5x in all three patient samples revealed consistently increased expression of RCC-associated genes (including *VEGFA, CA9, EGLN3*, etc.) in tumor cultures with concomitant downregulation of normal tubule-associated transcripts (e.g. *CDH1, ANPEP*) (**Supplemental Figure 2A**). Some tubule-derived genes did not change significantly in the RCC samples (e.g. *MME*). For subsequent analyses, we chose to anchor on genes that were expressed in our primary tumor cultures since the TCGA RNA-seq dataset is derived from whole kidney and tumor tissue and contains transcripts derived from non-tumor and non-tubule cell types (e.g. circulating immune cells, stromal cells, endothelial cells). Of genes that were expressed at a minimum threshold (FPKM≥1) in our samples, 1,072 genes were upregulated and 1,207 genes were downregulated across all three patient tumor samples compared to their respective tubule controls. Gene ontology analysis identified pathways characteristically dysregulated in RCC, such as genes related to the hypoxic response (e.g. *VEGFA*), organic ion transport (e.g. *CA9*) and lipid metabolism (e.g. *FABP6*), which were enriched in the upregulated gene set. Genes related to cell cycle regulation (e.g. *AURKA, TOP2B*) and chromatin organization (e.g. *HMGA1*) were consistently transcriptionally downregulated (**Supplemental Figure 2B**). Thus, the gene expression landscapes of our primary cultures are concordant across patient samples and recapitulate key transcriptional signatures of RCC.

**Supplemental Figure 2.**
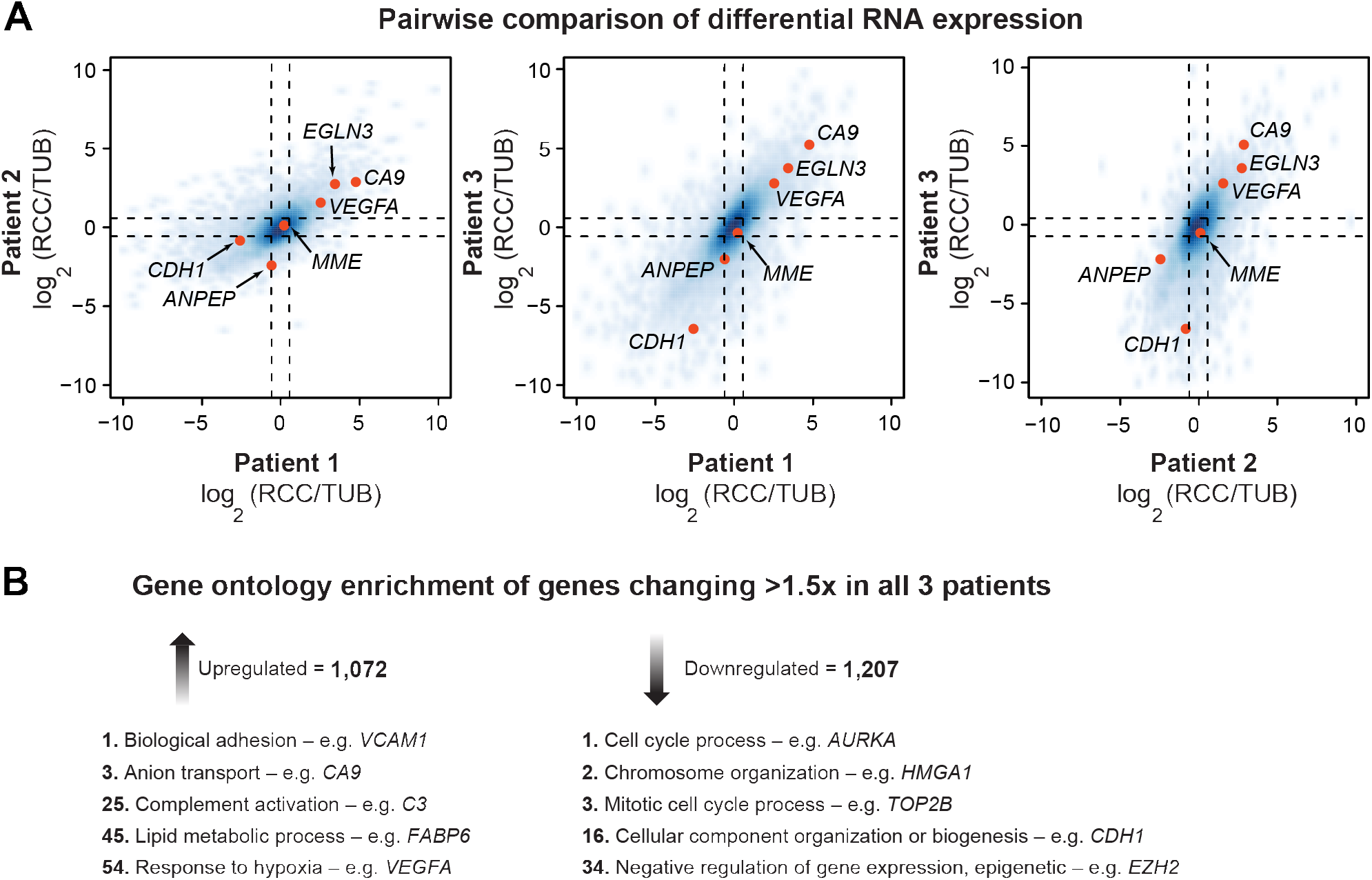
Individual renal cell carcinomas exhibit highly concordant RNA landscapes. (A) *Consistent patterns of gene expression among patients*. Comparison of expression fold change of genes reveals largely consistent patterns of gene expression among patient tumor samples. Genes that typify the HIF transcriptional response (e.g. CA9, *VEGFA, EGLN3*) are upregulated and some genes associated with normal tubular function (e.g. *CDH1, ANPEP*) are downregulated in all three tumor samples compared to their normal tubule controls. (B) *Gene ontology enrichment*. Ranked list GOrilla enrichment analysis (rank in boldface) identifies both canonical (e.g. hypoxia response, lipid metabolic process, chromatin organization) and unexpected gene ontologies (e.g. complement activation) that are differentially regulated in renal cell carcinoma compared to tubules.

### Concordant tumor regulatory landscapes expose transcription factor drivers of RCC

Chromatin accessibility profiling methodologies such as DNase-seq uniquely provide insight into the transcription factors drivers of oncogenesis (Stergachis et al. 2013). Since HIF is canonically dysregulated in RCC, we next explored its role and that of other transcription factors (TFs) in driving the chromatin accessibility changes we observed in the regulatory landscapes of the patients’ tumor samples. Even though most (>93%) HIF binding sites coincide with DHSs, ~70% of these DHSs show no significant change in accessibility between tubule and RCC (**Figure 3A**). Even the HIF-bound DHSs that showed significant accessibility changes in one tumor-normal pair often did not show differential DHS accessibility in the other patient samples (**Figure 3B**). This suggested that HIF alone does not broadly reprogram the regulatory landscape of RCC, but did not exclude the possibility that it may regulate other TFs that contribute to the process of malignant transformation. 213/776 of the TFs that are upregulated (≥1.5x) in at least one patient RCC-tubule pair have a HIF-occupied DHS within 250kb of their transcription start site (TSS) (**Figure 3C**). A subset of these 213 TFs shows evidence of restricted transcriptional induction in RCC compared to multiple somatic tumors for which matched normal tissues are available for comparison in the TCGA expression data (**Figure 3D**). Since the presence of a HIF-bound DHS near an induced TF gene does not conclusively demonstrate regulation of that gene by HIF, the TF gene subset that is induced in the TCGA data is more likely to contain TFs truly subject to HIF regulation in RCC. Alternatively, the fact that only a subset of the putative HIF-regulated TFs in our primary culture system shows selective expression in the TCGA RCC RNA-seq data may reflect the contaminating effect of non-tumor cell types in TCGA samples that can obscure small changes in transcription factor genes that are typically expressed at low levels.

**Figure 3.**
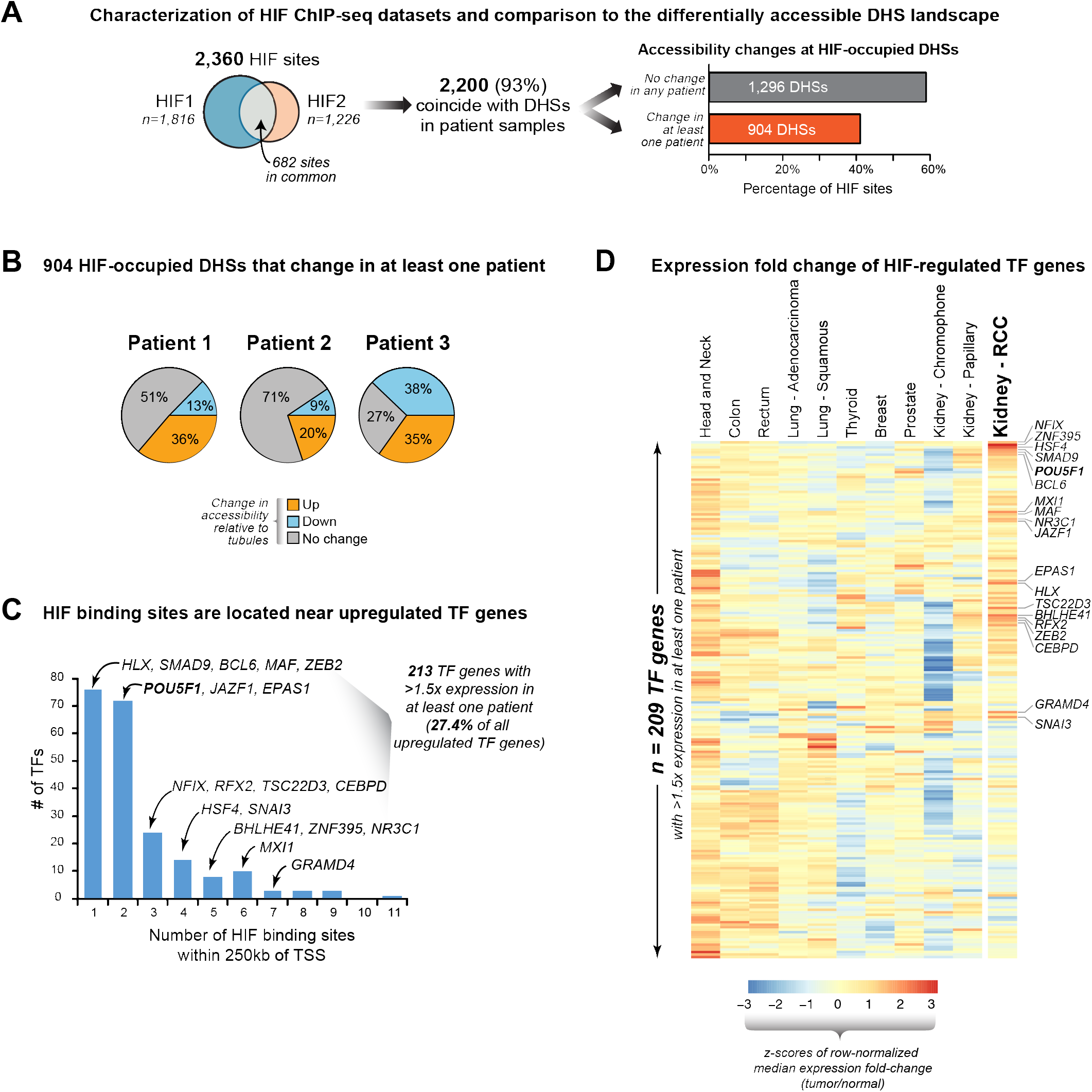
Concordant tumor regulatory landscapes expose key transcription factor drivers of RCC. (A) *HIF-binding only accounts for a small proportion of the differentially accessible RCC regulatory landscape*. ChIP-seq datasets for HIF1A and HIF2A show substantial overlap with each other and most of these peaks coincide with a DHS in the tubule and/or RCC DNase-seq datasets. Most HIF peaks in DHS map to non-changing/constitutive DHS in the tubules and RCC. (B) *Differentially accessible HIF-bound DHS show different patterns of accessibility across patient samples*. Of the 904 HIF peaks that map to differentially accessible DHS in at least one patient sample, most do not show significant change across the other patients’ samples. (C) *Transcription factors with changing expression located near HIF binding sites*. The expression levels of 213 transcription factors change by >1.5x in at least one patient sample and exhibit at least one HIF bound DHS within 250kb of their transcription start site (TSS). Many of these contain numerous HIF binding sites in proximity to their TSS, including transcription factors linked to renal cell carcinoma susceptibility (*ZEB2, BHLHE41*) and *POU5F1*. (D) *Selective expression of transcription factors in cancer*. The transcription factors that are expressed (FPKM>1) and changing by at least 1.5-fold in any of the three patient samples (from panel C) are examined for differential expression in a wide range of tumors that have matched normal tissues available in TCGA RNA-seq expression dataset (209 transcription factors are depicted; 4 factors are not mapped in the TCGA RNA-seq data). Transcription factors with RCC-selective increased expression are highlighted (e.g. *HSF4, BHLHE41, ZEB2, POU5F1*, etc.).

To uncover the identities of the TFs that are likely to be driving the regulatory program of RCC, we determined the relative enrichment of TF recognition sequences within the shared set of differential DHSs (discussed above) compared to a background of static DHSs. AP-1, ETS and E-box family recognition sequences were significantly enriched in DHSs with decreased accessibility in RCC (**Figure 4A**). Motifs for basic helix-loop-helix (bHLH) family transcription factors (which include *MYC, HIF* and *BHLHE41*) were enriched in DHSs that do not change their accessibility in RCC. Recognition sequences for several TF families (including homeodomain, nuclear receptor and HNF1/POU) were enriched in DHSs with increased accessibility in RCC.

Since several TF family members can recognize the same DNA binding recognition sequence, we next asked if the differential TF gene expression levels between tubules and RCC could help identify the specific family members that were contributing to the observed motif enrichment in the regulatory landscape. This analysis revealed that for the POU family transcription factors, only the stem cell related factor *POU5F1* (also known as OCT4) is consistently expressed and upregulated in RCC compared to tubules (**Figure 4B**). *POU5F1* and some of the transcription factors which are associated with genetic risk for RCC and whose binding sequences are enriched in differentially accessible DHSs (e.g. *BHLHE41*) show evidence of regulation by HIF (**Figure 3C**). *POU5F1* is normally expressed only in stem cells and germ cell-derived tumors but in the larger TCGA data set, it shows strikingly selective induction in RCC and papillary kidney cancer (both derived from proximal tubule cells) compared to normal kidney tissue (**Figure 4C**). Other known cellular reprogramming transcription factor genes, namely *SOX2, KLF4* and *NANOG*, are not induced in RCC (*data not shown*).

Taken together, these results suggest that instead of driving large-scale changes in chromatin accessibility by itself, HIF may have a broader impact on the regulatory landscape of RCC by activating other transcription factors. We sought to corroborate this notion by closer examination of the role of HIF in the regulation of *POU5F1*.

**Figure 4.**
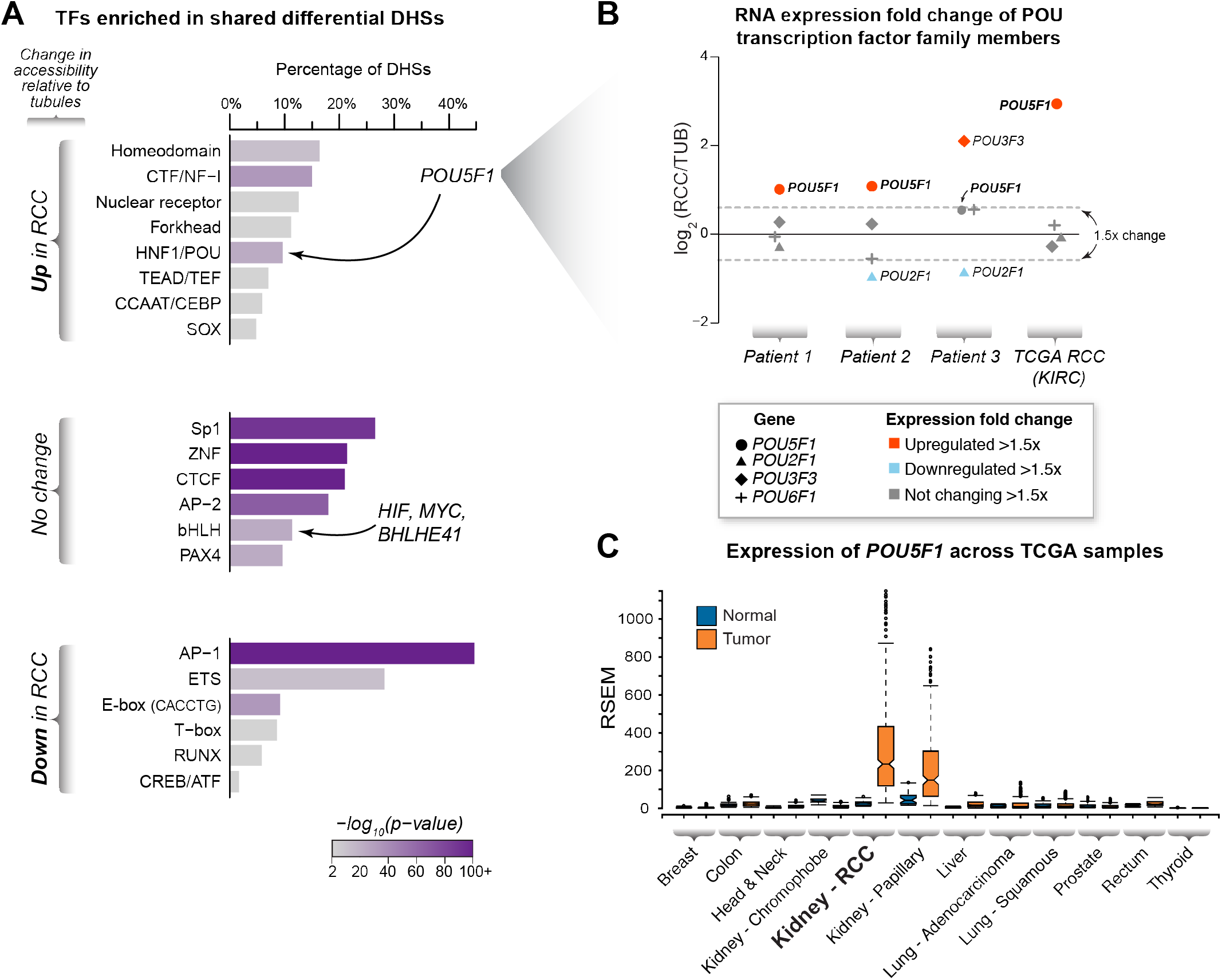
Correlation of DNA binding motif enrichments with gene expression identifies enrichment for *POU5F1* in RCC. (A) *Transcription factor enrichment*. Examination of differentially accessible or non-changing DHSs reveals different classes of transcription factors whose DNA binding recognition sequences are enriched in each category. The motif families containing transcription factors with genetic evidence linked to renal cell carcinoma susceptibility (i.e. *MYC, BHLHE41, ZEB2* and *HIF*) and the stem cell related transcription factor *POU5F1* (OCT4) are indicated. (B) *Examination of RNA expression identifies candidate POU-family transcription factors driving motif enrichments in the DHS landscape*. Since multiple transcription factors within the POU family share redundant DNA binding motifs, examination of transcription factor expression patterns may identify specific family members that are driving motif enrichment signatures. Examination of the differential gene expression patterns of these family members in RCC vs. tubules in our primary cultures and in the TCGA RNA-seq dataset reveals upregulation of *POU5F1* in RCC. (C) *Expression of POU5F1 in diverse somatic tumors*. The mRNA expression levels of the stem cell related transcription factor *POU5F1* (OCT4) in several non-germ cell tumors is compared to their matched normal tissue controls. The ends of the bar plots represent the 25^th^ and 75^th^ quartiles with whiskers representing 1.5x inter-quartile range (10% outlier trim applied).

### Expression of a novel POU5F1 transcript in RCC from a human- and kidney-specific promoter

Close examination of the chromatin accessibility and RNA-seq data from our three patients revealed a long, intergenic stretch of RNA transcription starting from a DHS and leading into (and on the same strand as) the annotated *POU5F1* transcripts (**Figure 5**; strand-specific signal not shown). This regulatory element, ~16 kb upstream of the *POU5F1* TSS used in embryonic stem (ES) cells, was distinct from the well-characterized distal and proximal enhancers that regulate *POU5F1* in ES cells (Nordhoff et al. 2001). Furthermore, this DHS was only present in adult kidney tubule- and RCC-derived cells/cell lines and was not detected in ES cells, fetal kidney tissues or many other diverse cell types (**Supplemental Figure 3**).

**Figure 5.**
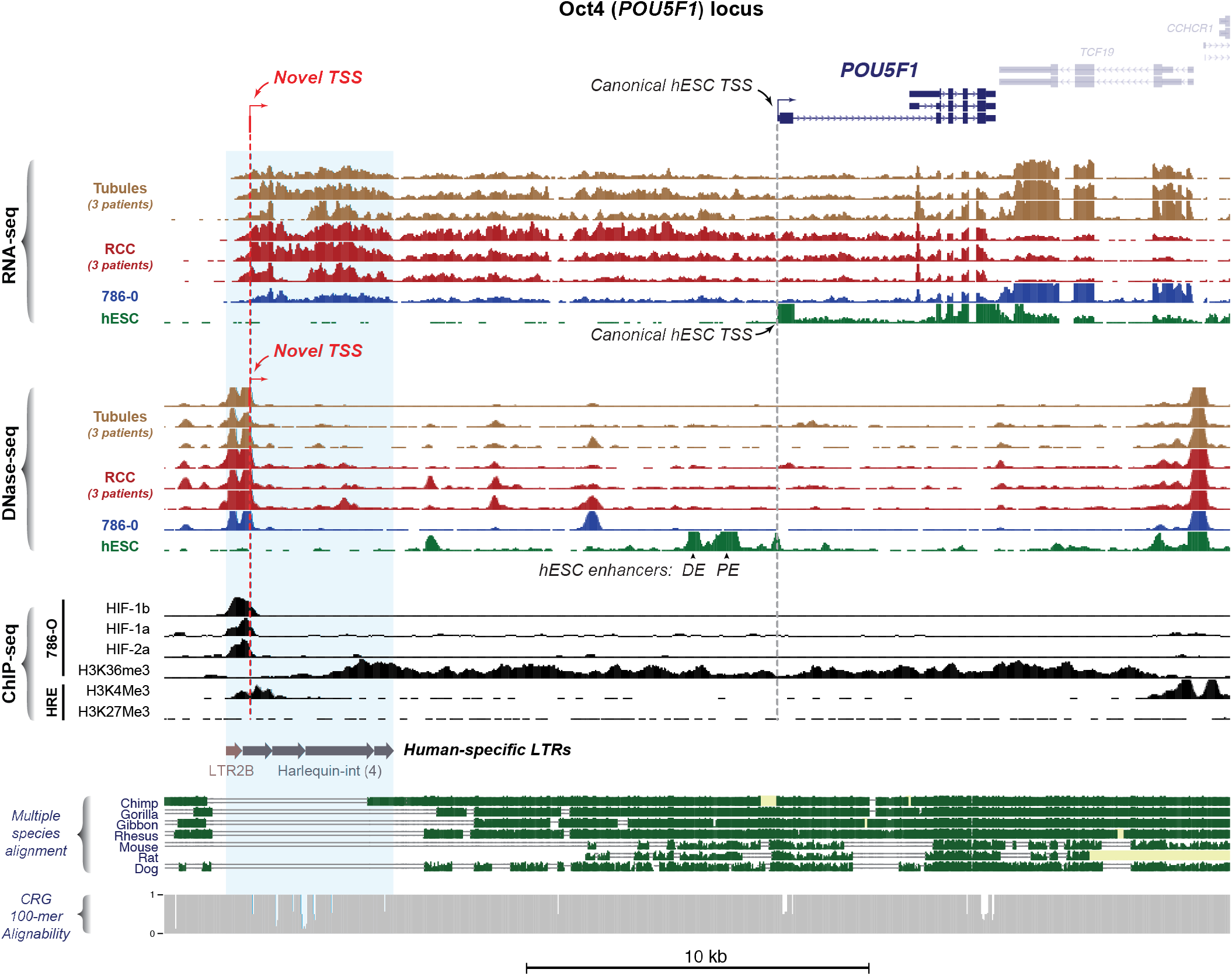
A novel human-specific promoter drives *POU5F1* expression in RCC. *Overview of the POU5F1 genomic locus (hg19 chr6:31,125,253-31,156,354*). RNA-seq tracks for the primary patient samples and the RCC cell line 786-O reveal a novel transcript originating from a DHS ~16kb upstream of the known ES cell transcription start site. ChIP-seq reveals binding of HIF components (HIF1 α, HIF2α, HIF1β) to this DHS with evidence of histone modification typical of active transcription across the entire transcript (H3K36Me3). This DHS is also associated with histone modifications characteristic of an active promoter, i.e. positioned nucleosomes marked by H3K4Me3 and depletion of the repressive H3K27Me3 mark. Examination of sequence conservation shows that this novel promoter lies within a complex tandem long terminal repeat element that is unique to humans.

**Supplemental Figure 3.**
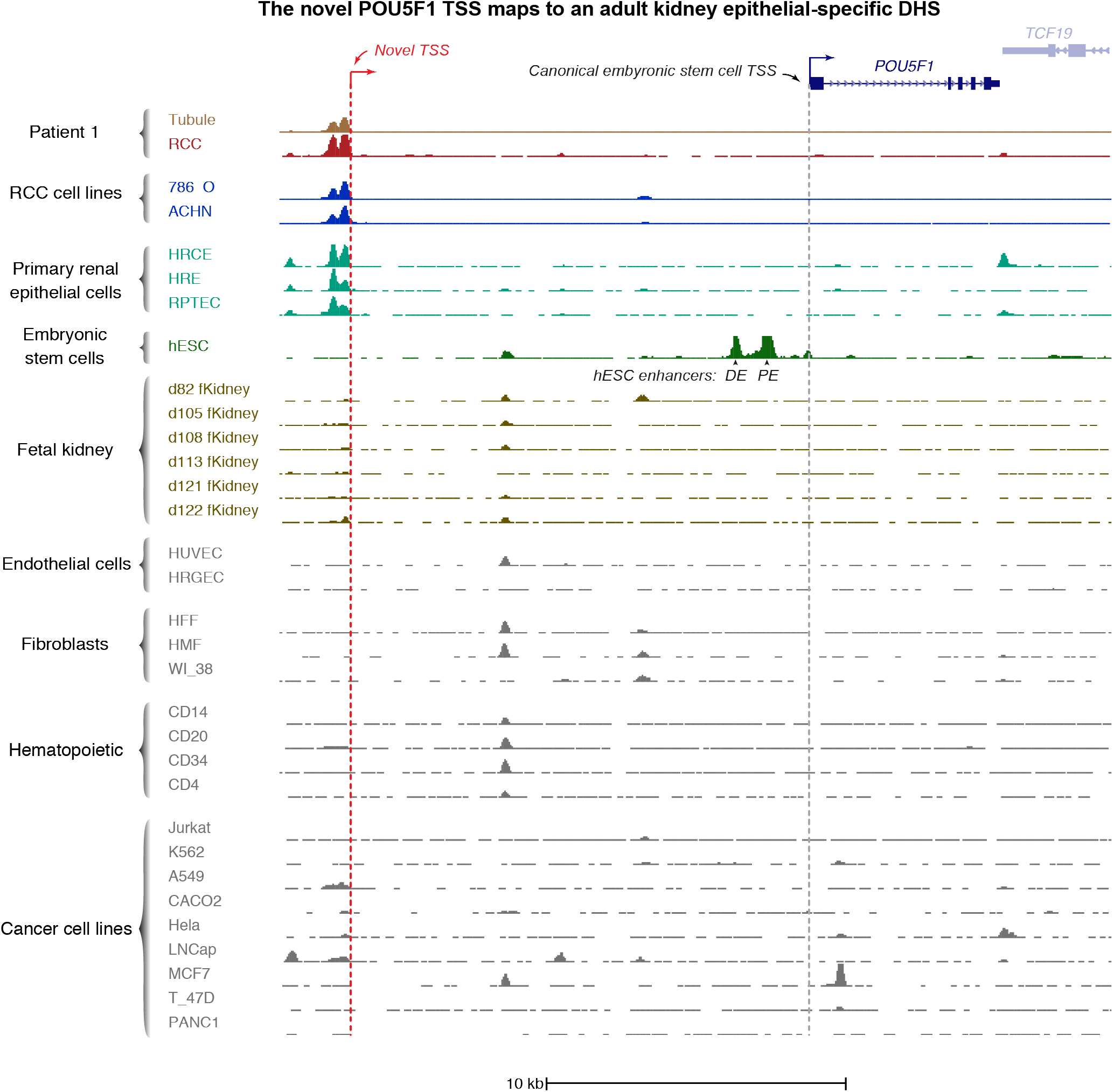
An adult kidney-specific DHS encodes a novel promoter for *POU5F1*. The novel transcription start site for *POU5F1* in RCC maps to a DHS that is only present in adult kidney derived tubules, primary cultures or tumors. This DHS is not present in fetal kidney, embryonic stem cells, non-epithelial kidney cells (e.g. glomerular endothelial cells, HRGEC) or a variety of ontologically diverse cells.

FANTOM5 data suggest that this kidney-specific DHS acts as a promoter: it coincides with H3K4me3, which marks active promoters; lacks H3K4me27; and demarcates H3K36me3 signal, a mark associated with transcription elongation, that extends into annotated *POU5F1* transcripts (Andersson et al. 2014; FANTOM Consortium and the RIKEN PMI and CLST (DGT) et al. 2014). We sought to determine whether novel transcripts of *POU5F1* were generated from the -16kb DHS in RCC. Knowing that the expression of *POU5F1* may be confounded by that of its pseudogene, *POU5F1B* (Takeda et al. 1992; Liedtke et al. 2007), we examined chromatin accessibility and gene expression at the *POU5F1B* pseudogene locus in our samples, and did not detect significant amounts of either (**Supplemental Figure 4**). We then proceeded to unambiguously determine if the putative promoter initiated transcription of a novel *POU5F1* isoform. To do this, we performed 5’-RACE on cDNA isolated from the VHL-null 786-O RCC cell line and sequenced the resulting products (**Figure 6A**). This captured a new transcription start site for *POU5F1* originating within the -16kb DHS. Several exon combinations were observed suggesting a complex mixture of isoforms expressed in 786-O cells.

Critically, the -16kb DHS coincided with strong HIF1α and HIF2α ChIP-seq signal in the 786-O cell line, suggesting that HIF is bound to this promoter element in RCC. We note that this HIF site is encoded by long-terminal repeat (LTR) elements of the Harlequin-int and LTR2B subfamilies of ERV1 endogenous retroviruses. This repeat configuration appears to represent an evolutionarily recent insertion into the human genome as it is not conserved among higher primates or other mammals (**Figure 5**). Good CRG alignability (Derrien et al. 2012) at this composite LTR reduced the possibility that degeneracy of viral repeat elements may confound locus-specific mapping of short-read sequences.

Finally, we asked if the canonical and novel isoforms of *POU5F1* exhibited dependence on VHL protein (stably reintroduced into the 786-O cell line) and/or hypoxia using isoform specific RT-PCR primers (**Figure 6A**). Reintroduction of VHL protein into 786-O cells cultured in normoxia strongly suppressed expression of both canonical and novel *POU5F1* transcripts (**Figure 6B**). The presence of VHL protein also resulted in significant induction of canonical and novel *POU5F1* transcripts when the 786-O+VHL cells were cultured in hypoxia (**Figure 6B**). These transcripts did not change appreciably when 786-O cells were shifted from normoxia to hypoxia, consistent with already maximal HIF-signaling in this *VHL*-null cell line.

Taken together, these results establish the presence of a HIF-responsive, kidney-specific promoter element that initiates expression of a novel transcript of *POU5F1* in RCC and originated at this locus by insertion of endogenous retrovirus elements within the human lineage.

**Supplemental Figure 4.**
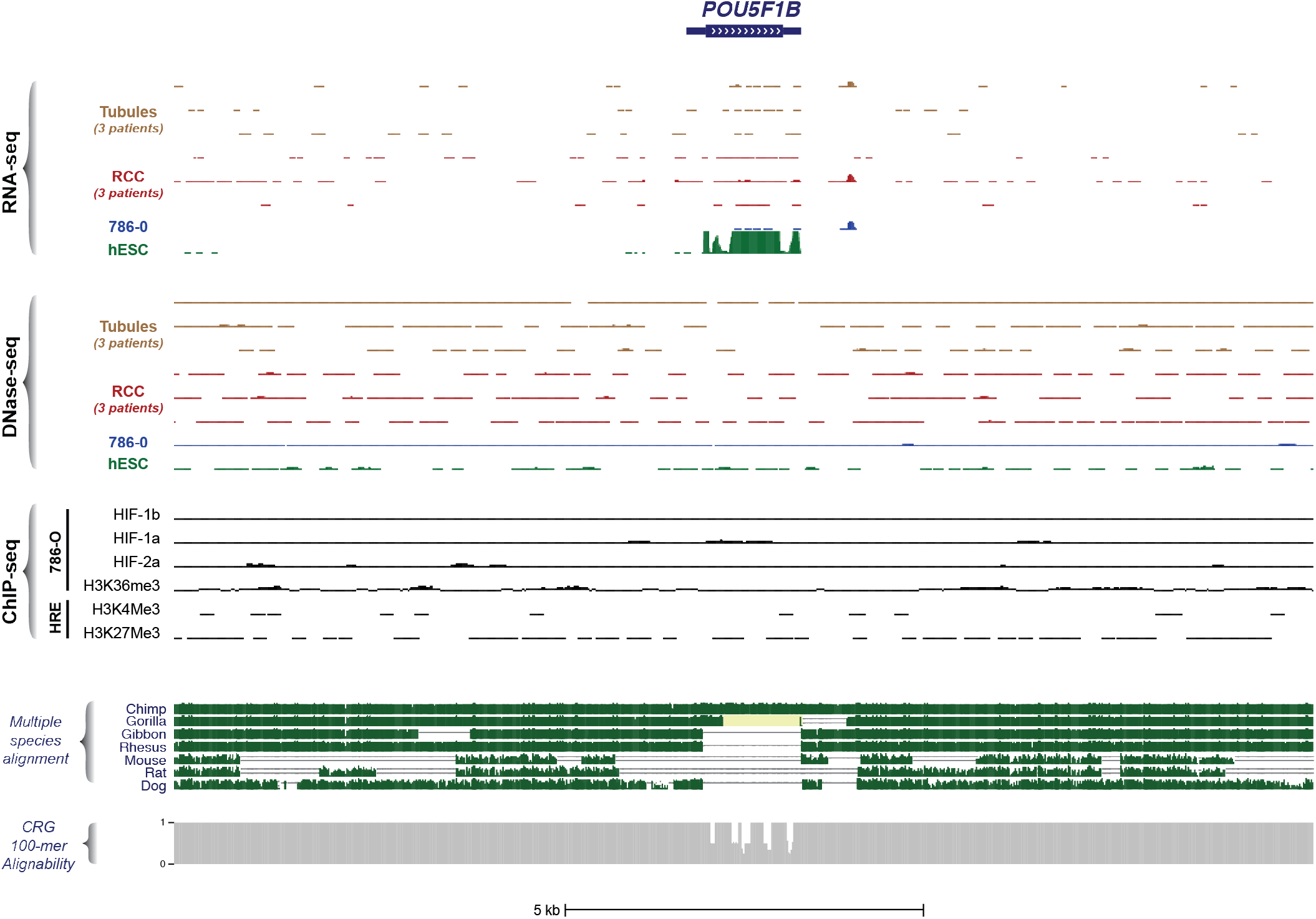
Overview of the *POU5F1B* genomic locus (hg19 chr8:128,420,724-128,436,573) *POU5F1B* is expressed in human ESCs, but not in the primary tubule and RCC cultures described in this study. There are no DHSs in this genomic interval and there is negligible binding of HIF components. Histone modifications typical of active transcription are also not present.

**Figure 6.**
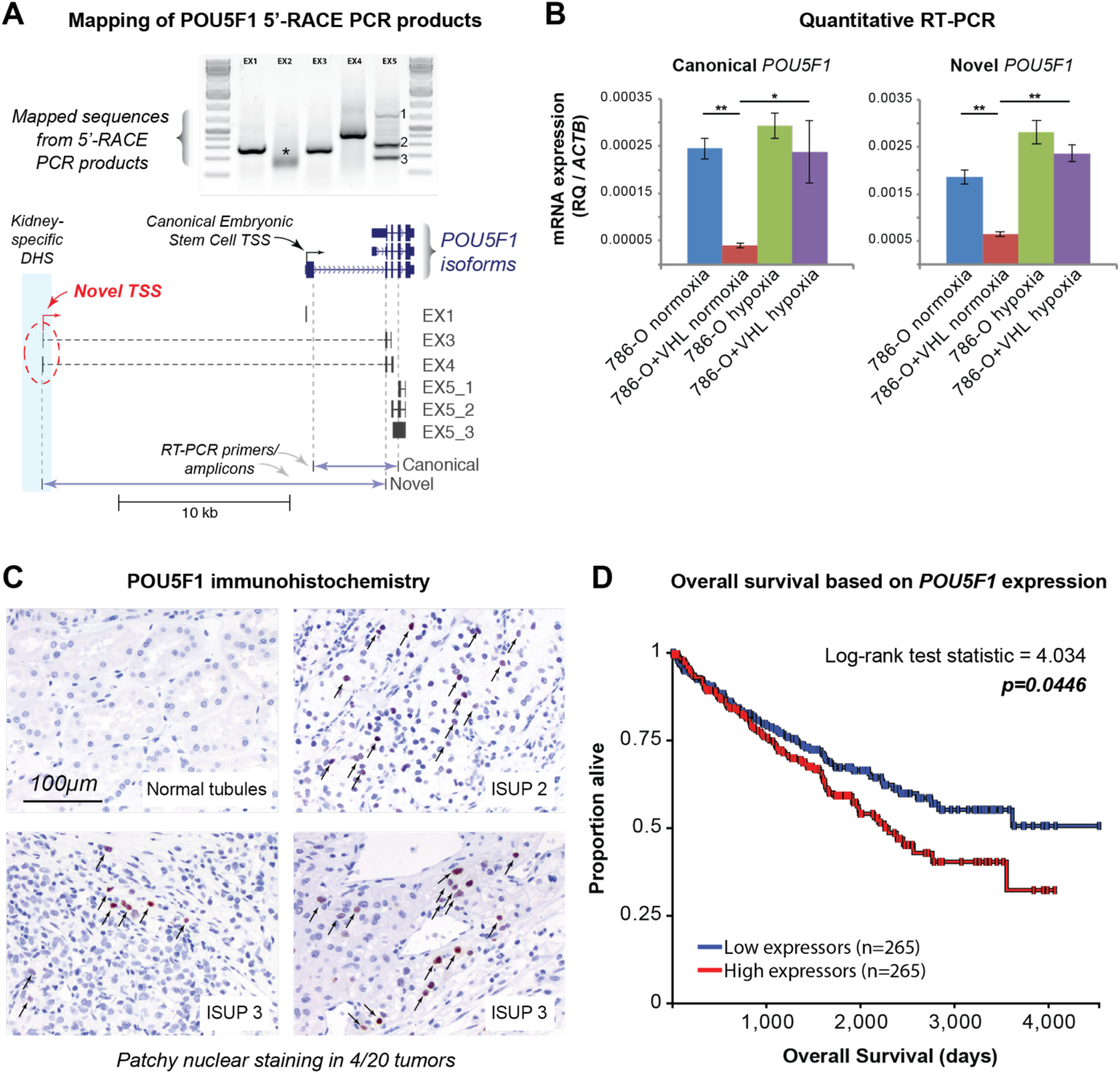
The novel transcript for *POU5F1* exhibits HIF dependence and *POU5F1* expression levels correlate with patient survival. (A) *Schematic mapping of POU5F1 5’-RACE PCR products*. 5’-RACE performed on 786-O RNA captures a transcription start site originating in the novel DHS therefore defining a novel isoform of *POU5F1*. Reverse primers in known *POU5F1* exons (e.g. EX1 = reverse primer in exon 1) were used to amplify the 5’-ends of the cDNA molecule captured by 5’-RACE and sequence mapped to the genome. The exon 2 primer (*) failed to yield mappable sequence. The exon 5 primer yielded 3 different products (EX5-1, EX5-2, EX5-3). The location of PCR primers to detect the canonical and novel *POU5F1* transcript variants are indicated. (B) *Canonical and novel POU5F1 transcripts exhibit HIF-dependence*. RT-PCR primers were used to quantify the canonical and novel POU5F1 transcripts in 786-O cells and 786-O cell stably transduced with VHL (786-O+VHL) cultured in normoxia or hypoxia (2% O_2_) for 24 hours. Expression levels (relative quantification, RQ) were calculated using the β-actin housekeeping gene (*ACTB*). Reintroduction of VHL protein into 786-O cells suppresses expression of both *POU5F1* transcripts. Exposing 786-O+VHL cells to hypoxia induces both *POU5F1* transcripts. Error bars indicate standard deviations of three experimental replicates. * p<0.05, ** p<0.005. (C) *Immunohistochemistry of POU5F1 protein in renal cell carcinoma samples*. POU5F1 (OCT4) immunohistochemistry was performed on RCC samples from 20 patients (5 from each of ISUP grades 1-4) and showed patchy nuclear positivity (arrows) in a single random sample from 4 patients. No nuclear staining was seen in any of the matched normal renal parenchyma from the same patients. (D) *Overall survival as a function of POU5F1 expression in TCGA*. Patients with *POU5F1* expression data from TCGA (KIRC) were evenly divided into two groups split at the median expression level (233 RSEM normalized) and Kaplan-Meier curves for overall survival were plotted using the UCSC Xena browser tool.

### POU5F1 expression in RCC correlates with overall survival

Next, we sought to evaluate if induced *POU5F1* transcription led to increased protein levels in human RCC specimens. For these experiments, we utilized an antibody recognizing a C-terminal epitope of POU5F1 (OCT4) that is expected to be represented in both the canonical and novel isoforms of *POU5F1*. Preliminary experiments using a tissue microarray with 102 cases of localized RCC and 25 cases of advanced stage/metastatic RCC did not reveal significant POU5F1 (OCT4) expression in the tumor cells (data not shown). However, since the tissue cores for each individual tumor in the array are very small and may not be representative of the often large and heterogeneous RCC tumors (Gerlinger et al. 2012; 2014), we decided to test POU5F1 (OCT4) expression in larger tissue sections from 20 different patient tumors alongside their matched normal kidney controls. In 4 out of 20 RCC tissue sections, patchy nuclear POU5F1 (OCT4) protein expression was detectable (Chi-squared p-value=0.035, **Figure 6C**). We did not observe POU5F1 (OCT4) expression in any of the normal kidney tissue sections examined. Therefore, even though *POU5F1* transcript induction appears to be a consistent feature of RCC, POU5F1 (OCT4) protein is less frequently detected which may reflect focal or patchy expression in these large tumors. Lastly, we examined *POU5F1* expression in the TCGA data set as a function of clinical staging parameters. The expression of *POU5F1* did not correlate with metastasis status (**Supplemental Figure 5A**), but was positively correlated with pathologic tumor stage, with higher stage tumors exhibiting greater expression of *POU5F1* (**Supplemental Figure 5B**). Strikingly, patients with high expression of *POU5F1* exhibited lower overall survival compared to patients with lower expression levels (**Figure 6D**). These results demonstrate that POU5F1 (OCT4) protein can be expressed in a patchy fashion in RCC tumors and that *POU5F1* expression levels can predict overall survival in patients with RCC.

**Supplemental Figure 5.**
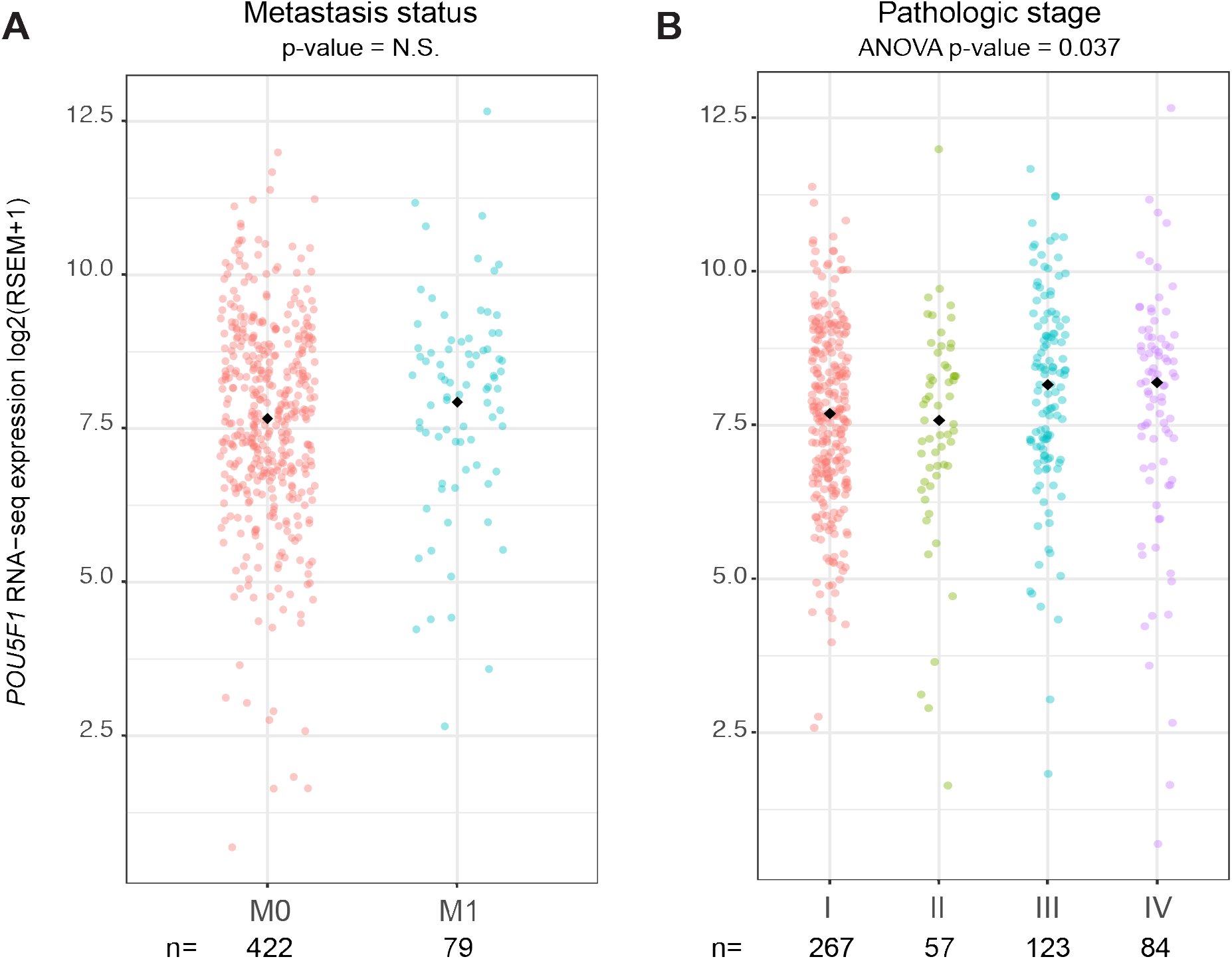
Expression of *POU5F1* in RCC as a function of known metastasis status and pathologic stage. Black diamond indicates mean value for the indicated subgroup in both panels. (A) Expression of *POU5F1* as a function of known metastasis status. (B) Expression of *POU5F1* as a function of pathologic stage at the time of diagnosis. P-value = 0.037 by 1-factor ANOVA.

## Discussion

Even for a well-studied tumor such as RCC, there is a notable deficit in the understanding of genome dysregulation that drives oncogenesis. Here we demonstrate that while each patient’s tumor can exhibit its own unique epigenomic signature, subtraction of the genotype-matched cell-of-origin baseline and comparison across individuals can identify the core regulatory landscape of cancer. Using high-resolution epigenomic mapping on primary tumors and matched normal cells from three patients, we identified multiple transcription factors with differential expression patterns and significant DNA binding motif enrichments that likely contribute to the tumor phenotype. Transcription factors that drive genome dysregulation in RCC have hitherto only been explored in piecemeal fashion. Besides the HIFs, other sequence-specific factors have been implicated individually in various aspects of RCC biology including PAX2 (Daniel et al. 2001; Doberstein et al. 2011; Gnarra and Dressler 1995; Luu et al. 2009), PAX8 (Hu et al. 2012; Laury et al. 2011; Tong et al. 2011), CEBPβ (Oya et al. 2003), NRF2 (Kinch et al. 2011; Ooi et al. 2013), FOXO (Cho and Mier 2012; Gan et al. 2010; Wu et al. 2013a), STAT3 (Bill et al. 2012; Horiguchi et al. 2002; 2010; Jung et al. 2005; Li et al. 2008; Xin et al. 2009; 2011), FOXM1 (Wu et al. 2013b; Xue et al. 2012), OCT4 (Bussolati et al. 2008; Smith et al. 2011), P53 (Oda et al. 1995; Reiter et al. 1993; Torigoe et al. 1992; Uhlman et al. 1994), TCF21 (Ye et al. 2012; Zhang et al. 2012), HCF1 (Peña-Llopis et al. 2012), HNF1/2 (Anastasiadis et al. 1999; Rebouissou et al. 2005; Sel et al. 1996) and most recently BHLHE41 (Bigot et al. 2016; Grampp et al. 2017) and ZNF395 (Rhie et al. 2016; Zhao et al. 2016). Here, we show that many of these transcription factors may in fact be regulated by HIF and appear to influence the regulatory landscape in RCC.

One transcription factor that is consistently upregulated in RCC and influences its regulatory landscape is the stem cell factor *POU5F1 (OCT4). POU5F1*, together with *KLF4, SOX2* and *NANOG* (which are not expressed in RCC) is well known for its ability to reprogram somatic cells into pluripotent stem cells (Park et al. 2008). Hypoxia is a known stimulant of *POU5F1* expression in embryonic stem and cancer cells (Ezashi et al. 2005; Westfall et al. 2008; Forristal et al. 2009; Mathieu et al. 2011) and can even reprogram committed cells into a pluripotent state (Mathieu et al. 2013; 2014). Our examination of the *POU5F1* genomic locus identified a novel adult kidney-selective and hypoxia/HIF-responsive promoter that produces a previously undescribed transcript isoform for *POU5F1* in RCC. We also show that this novel promoter lies within a tandem, human-specific LTR element. Repeat elements such as LTRs are enriched in primate-specific regulatory elements (Jacques et al. 2013) and are known to influence transcription factor regulatory networks (Bourque et al. 2008). Therefore, these findings suggest that regulation of *POU5F1* expression in adult human kidney epithelial cells, either in response to hypoxia or constitutive HIF activation as occurs in RCC, may differ significantly from human embryonic stem cells and mouse tissues and requires separate, context-specific study.

Activation of stem cell-like epigenetic and transcriptional programs are associated with malignant transformation, though clear cell RCC appears to behave differently than other tumor types (Malta et al. 2018). Compared to other somatic cell types (Park et al. 2008), human kidney proximal tubule cells appear to have a lower barrier to reprogramming to pluripotency as they require only *SOX2* and *OCT4* expression (Montserrat et al. 2012). Since *VHL* inactivation and constitutive HIF stabilization appear to be early events in sporadic RCC (Mitchell et al. 2018; Turajlic et al. 2018), it will be important to determine if this genetic lesion alone is sufficient to induce *POU5F1* expression in kidney proximal tubule cells. Interestingly, we found that the level of *POU5F1* expression appears to predict patient survival even though only a subset of tumor cells appear to produce OCT4 protein, perhaps marking RCC cancer stem cells (Bussolati et al., 2008). In particular, given the documented intratumoral heterogeneity and divergence of metastatic RCC clones (Gerlinger et al. 2012; 2014), it will be necessary to compare the epigenomic profiles of those samples with that of the primary tumor from multiple patients. Taken together, understanding the mechanisms that activate *POU5F1* expression from this promoter in adult kidney epithelial cells and its effects on cellular transformation and clinical behavior will be intriguing topics for future studies.

The data generated and described here are freely available to provide a reference map upon which future functional genomic studies on RCC can be constructed and interpreted. Overall, our approach demonstrates the power of epigenomic analysis focused on small numbers of pure primary tumor and matched normal cell-of-origin cultures which can provide a clarifying lens through which to interpret inherently noisier large tumor-sequencing datasets. This general framework can reveal unanticipated insights into tumor biology and is readily applicable to other cancers in which tumor cells and matched normal cells-of-origin are available.

## Methods

### Patient tissue sample procurement and primary cell culture

Malignant and normal kidney tissues were obtained from patients undergoing radical nephrectomy for clear cell renal cell carcinoma with informed consent for DNA sequencing obtained prior to the surgery. The study (#1297) and consent forms were approved by the University of Washington’s IRB. Patient 1’s cultures were derived from an 80-year-old woman; Patient 2’s cultures were derived from a 62-year-old man and Patient 3’s cultures were obtained from a 63-year-old man. At the time of surgery, all patients presented with localized disease. Approximately 1cm^3^ portions of tumor (from a central, non-necrotic location) and uninvolved kidney cortex (usually from the pole furthest from the tumor mass) were harvested and transported in RPMI medium on ice. These tissues were then minced with a sterilized razor blade and the resulting fragments were placed in 20mls of pre-warmed RPMI medium (without serum) supplemented with Accutase (Sigma, diluted 1:10), collagenase P (Roche, 100μg/ml) and trypsin/EDTA (Gibco, 0.25% solution diluted 1:10). The tissue fragments were digested at 37°C for 20 minutes with vigorous agitation. After digestion, the tissue fragments were spun down and macerated with a sterile plunger from a 5-ml syringe. These softened tissue fragments were then transferred into tissue culture flasks with pre-warmed culture medium (RPMI supplemented with 10% fetal bovine serum and ITS+ supplement, Corning). After 3-4 days (for tubule cultures) and 7-10 days (for RCC cultures), the tissue fragments were decanted and the adherent cells were fed with fresh medium. At this stage, primary tubule cells grew rapidly and had an epithelioid morphology, while primary RCC cells grew slowly, were larger and exhibited frequent cytoplasmic vacuoles typical of adenocarcinoma. Cells were sub-cultured 1:4 when they reached 80% confluence and used within two passages for all experiments.

### 786-O and ACHN cell culture

The VHL-null 786-O (CRL-1932) and VHL-wildtype ACHN (CRL-1611) renal cell carcinoma cell lines were obtained from ATCC. Cells were cultured in RPMI medium supplemented with 10% fetal bovine serum, non-essential amino acids, glutamine and penicillin/streptomycin. Cells were sub-cultured 1:10 when they reached 80% confluence using Accutase to disaggregate adherent cells.

### Processing of cell cultures for DNase-seq

Primary tubule and RCC cultures, 786-O and ACHN cells were subjected to DNase I treatment, small DNA fragment isolation and double-stranded library construction per published ENCODE protocols or a recently described low-input single-stranded library construction protocol (Gansauge and Meyer 2013; Snyder et al. 2016). Libraries were subjected to paired-end (2x36bp) sequencing. The majority of datasets used in this study were deemed of high quality (signal portion of tags, SPOT>0.4) (Thurman et al. 2012). See **Supplemental Table 1** for cell input, quality metrics and other sequencing metadata.

### Processing of cell cultures for RNA-seq

Disaggregated cells from primary tubule or renal cell carcinoma cultures, 786-O and ACHN cells were washed once in PBS and stabilized in RNALater (Ambion). Total RNA was extracted using a mirVana RNA isolation kit (Ambion). Illumina sequencer compatible libraries were constructed using a TruSeq Stranded Total RNA Library Prep Kit with Ribo-Zero Gold (Illumina) and subjected to paired-end (2x76bp) sequencing. See **Supplemental Table 1** for cell input, quality metrics and other sequencing metadata.

### Karyotyping of primary cell cultures

G-band karyotyping of the primary renal cell carcinoma cultures was performed by the University of Washington Cytogenetics and Genomics Laboratory in the Department of Laboratory Medicine.

### Assessing VHL status of primary cell cultures

Genomic DNA from 200,000 cells from each of the primary cultures was extracted using an ArchivePure DNA purification kit from 5Prime. Oligonucleotide primers covering exons 1-3 of the *VHL* gene (VHL_exon1_F1, GCGCGAAGACTACGGAGGTC; VHL_exon1_R1, CGTGCTATCGTCCCTGCT; VHL_exon2_F1, TCCCAAAGTGCTGGGATTAC; VHL_exon2_R1, TGGGCTTAATTTTTCAAGTGG; VHL_exon3_F1, TGTTGGCAAAGCCTCTTGTT; VHL_exon3_R1, AAGGAAGGAACCAGTCCTGT) were used to amplify genomic sequence using KAPA HiFi Taq polymerase (Kapa Biosystems). The resulting PCR products were separated on an agarose gel, purified and subjected to Sanger sequencing (EuroFins Scientific).

### 5’-RACE for novel POU5F1 transcripts

Total RNA was extracted from 7x10^6^ 786-O cells using the RNeasy Mini kit (QIAGEN cat #74104) according to manufacturer’s protocol. We then used 9 μg total RNA input for RLM-RACE (ThermoFisher Scientific First-Choice RLM-RACE, cat# AM1700), following the manufacturer’s “standard scale” 5’-RACE protocol, which ligates an adapter to the 5’ end of full-length, capped mRNA molecules. The primary PCR reaction was carried out using a common forward primer recognizing the 5’-RACE adapter and reverse primer located in each of the first five coding exons of POU5F1 (“R2” primers), using cycling conditions 94°C 3min, 35 cycles of 94°C 3min/60°C 30sec/72°C 2min, 72°C 7min. Of the 50μl primary PCR, 2μl was used for a secondary PCR with nested primers in the 5’-RACE adapter and within each of the five POU5F1 coding exons (“R1” primers), using the same cycling conditions as the primary PCR. Secondary PCRs were run on an agarose gel, the bands were excised and purified using a MinElute Gel Extraction kit (QIAGEN cat #28604) according to the manufacturer’s protocol, and were sequenced from both ends using Sanger sequencing.

### RT-PCR for canonical and novel POU5F1 transcripts

A clone of the VHL-null 786-O RCC cell line stably transduced with VHL (786-0+VHL) was originally obtained from Dr. William Kaelin’s laboratory (Yan et al. 2007). Approximately 200,000 786-O and 786-0+VHL cells were exposed in triplicate to hypoxia (2% O_2_) or normoxia for 24 hours. RNA was extracted using the RNeasy Plus Mini Kit (Qiagen, Valencia, CA), cDNA was synthesized using random hexamers and the Superscript IV First-Strand Synthesis Kit and was used to seed triplicate real-time PCR reactions using SYBR Green and standard cycling conditions for the Applied Biosystems 7900HT thermocycler. Primers were canonical *OCT4* (5’-GAGCAAAACCCGGAGGAGT-3’ and 5’-TTCTCTTTCGGGCCTGCAC-3’); novel *OCT4* (5’-GCTTGGCAAATTGCTCGAGTT-3’ and 5’-TGGAGTCCGGACATCTGAAAC-3’), and *ACTB* (5’-TCCCTGGAGAAGAGCTACG-3’ and 5’-GTAGTTTCGTGGATGCCACA-3’). A single peak was observed in the dissociation curve analysis for all genes and the sequence of the novel *OCT4* PCR product was confirmed by Sanger sequencing using the same primers. Cycle threshold (Ct) values were determined using Applied Biosystems Sequence Detection software. Relative quantification was calculated as 2^-delta Ct^, where delta Ct values were determined by subtracting the *ACTB* mean Ct values from the target gene Ct values.

### OCT4/POU5F1 immunohistochemistry

A tissue microarray (TMA) composed of cores of 102 cases of localized clear cell RCC, 25 cases of advanced/metastatic RCC, 62 cases of papillary RCC, 50 cases of chromophobe RCC/oncocytic neoplasms and 25 normal kidney controls was prepared with institutional IRB approval (study 9138). Twenty randomly selected RCC specimens (5 in each ISUP grade 1-4) were identified by a third-party honest broker, Northwest Biotrust at the University of Washington. One TMA section or a single section from each of the tumor mass and adjacent uninvolved kidney cortex were subjected to antigen retrieval with HIER ER1 buffer for 20 minutes (ER1= Epitope Retrieval Buffer 1, Citrate based pH 6.0 solution). Immunohistochemistry for OCT4/POU5F1 was performed using a 1:250 dilution of the OCT-3/4 (C-10) mouse monoclonal antibody (catalog # sc5279 from Santa Cruz Biotechnology).

**Supplemental Table 1.** Sample characteristics and sequencing metadata.

RNA-Seq
| GEO Accession | Patient |  | Sample ID | Gender | Age | Input cell |  | Total mapped reads (2x76bp) |
| --- | --- | --- | --- | --- | --- | --- | --- | --- |
|  | ID/cell line | Sample type |  |  |  | number | RIN |  |
| GSM3290917 | HIM13 | Human_kidney_tubule | DS33394 | F | 80 | 2,000,000 | 10 | 244,151,932 |
| GSM3290918 | HIM13 | Human_renal_cell_carcinoma | DS33395 | F | 80 | 500,000 | 9.9 | 257,438,823 |
| GSM3290919 | HIM15 | Human_kidney_tubule | DS37923 | M | 62 | 2,000,000 | 9.9 | 124,435,785 |
| GSM3290920 | HIM15 | Human_kidney_tubule | DS37924 | M | 62 | 2,000,000 | 9.9 | 104,505,925 |
| GSM3290921 | HIM15 | Human_renal_cell_carcinoma | DS37925 | M | 62 | 125,000 | 9.9 | 99,037,346 |
| GSM3290922 | HIM15 | Human_renal_cell_carcinoma | DS37926 | M | 62 | 125,000 | 9.8 | 147,534,033 |
| GSM3290923 | HIM23 | Human_kidney_tubule | DS40494 | M | 63 | 500,000 | 7.2 | 246,217,904 |
| GSM3290924 | HIM23 | Human_renal_cell_carcinoma | DS40496 | M | 63 | 500,000 | 9.1 | 113,565,853 |
| GSM3290925 | 786-O | Human_renal_cell_carcinoma | DS34766 | M | 58 | 4,750,000 | 9.8 | 187,589,844 |
| GSM3290926 | ACHN | Human_renal_cell_carcinoma | DS37193 | M | 22 | 2,250,000 | 7.4 | 85,933,914 |

DNase-seq
| GEO Accession | Patient |  | Sample ID | Gender | Age | Input cell |  | Total mapped reads (2x36bp) |
| --- | --- | --- | --- | --- | --- | --- | --- | --- |
|  | ID/cell line | Sample type |  |  |  | number | SPOT |  |
| GSM3291010 | HIM13 | Human_kidney_tubule | DS26689 | F | 80 | 9,000,000 | 0.5650 | 201,833,536 |
| GSM3291012 | HIM13 | Human_kidney_tubule | DS27824 | F | 80 | 9,000,000 | 0.4462 | 211,342,862 |
| GSM3291022 | HIM13 | Human_renal_cell_carcinoma | DS26693A | F | 80 | 5,000,000 | 0.5721 | 295,184,545 |
| GSM3291023 | HIM13 | Human_renal_cell_carcinoma | DS26692B | F | 80 | 5,000,000 | 0.4865 | 39,174,471 |
| GSM3291014 | HIM15 | Human_kidney_tubule | DS37969 | M | 62 | 2,000,000 | 0.3638 | 53,455,680 |
| GSM3291015 | HIM15 | Human_kidney_tubule | DS37971 | M | 62 | 2,000,000 | 0.4493 | 350,791,957 |
| GSM3291016 | HIM15 | Human_renal_cell_carcinoma | DS37973 | M | 62 | 300,000 | 0.4951 | 260,802,880 |
| GSM3291017 | HIM15 | Human_renal_cell_carcinoma | DS37974 | M | 62 | 300,000 | 0.5024 | 55,333,228 |
| GSM3291020 | HIM23 | Human_kidney_tubule | DS41160 | M | 63 | 80,000 | 0.3944 | 221,916,751 |
| GSM3291021 | HIM23 | Human_kidney_tubule | DS41396 | M | 63 | 80,000 | 0.5052 | 34,333,942 |
| GSM3291018 | HIM23 | Human_renal_cell_carcinoma | DS40508 | M | 63 | 80,000 | 0.2764 | 201,007,725 |
| GSM3291019 | HIM23 | Human_renal_cell_carcinoma | DS40509 | M | 62 | 80,000 | 0.2179 | 55,078,250 |
| GSM3291011 | 786-O | Human_renal_cell_carcinoma | DS27192 | M | 58 | 9,000,000 | 0.3238 | 214,308,625 |
| GSM3291013 | 786-O | Human_renal_cell_carcinoma | DS37199 | M | 58 | 100,000 | 0.3742 | 360,466,731 |
| GSM3291024 | ACHN | Human_renal_cell_carcinoma | DS24547A | M | 22 | 13,100,000 | 0.4583 | 261,250,750 |
| GSM3291025 | ACHN | Human_renal_cell_carcinoma | DS24471A | M | 22 | 3,000,000 | 0.4256 | 224,800,319 |

### DNase-seq data

Sequence reads from our DNase-seq libraries were subjected to an in-house uniform data processing pipeline, which we have used previously for ENCODE DNase-seq datasets (Thurman et al. 2012). Briefly, read pairs passing quality filters are trimmed of adapter sequences and aligned to the reference human genome (GRCh37/hg19) using BWA (Li and Durbin 2009). Genomic regions with a significant enrichment of DNase I cleavages were identified using our hotspot algorithm (Thurman et al. 2012) and were further refined to fixed-width, 150-base-pair regions (“peaks”) containing the highest cleavage density (referred to as DNase I hypersensitive sites, DHSs). Hotspot (FDR 1%) and peak calling were performed using both full-depth and uniformly sub-sampled (to 3.8 x 10^7^ aligned read pairs) data. Also see **Supplemental Table 1**.

### HIF ChIP-seq data

We downloaded sequence reads from ChIP-seq experiments for HIF-1α, HIF-2 α and HIF-1 β (Salama et al. 2015) from GEO (accession GSE67237), aligned them to the reference human genome (GRCh37/hg19) using BWA and identified peak summit locations using the macs2 algorithm (Zhang et al. 2008).

### RNA-seq data

RNA-seq libraries were aligned to the reference human genome (GRCh37/hg19) using TopHat 2.0.13 (Trapnell et al. 2009) and assigned to transcript models (GENCODE v19 basic set) using Cufflinks 2.1.1 (Trapnell et al. 2013). Also see **Supplemental Table 1**. Processed RNAseqV2 expression tables from TCGA Research Network (http://cancergenome.nih.gov/) were downloaded for frozen tissue samples from organ sites with matched normal and tumor tissues available for comparison. Patient annotations (e.g. tumor stage, metastasis status) for TCGA patient samples were obtained using the UCSC Xena browser tool (Goldman et al. 2018).

### General data processing

Data analyses were carried out using custom R scripts that utilized Bioconductor (http://www.bioconductor.org) packages for analyzing high-throughput sequencing data, custom Python scripts, and the BEDOPS (Neph et al. 2012) suite of tools, as well the publicly available tools GoRILLA (Eden et al. 2009) and GREAT (McLean et al. 2010) where indicated.

### Generation of DHS master list

To facilitate comparisons at the same genomic locus across multiple samples, we created a “master list” of non-overlapping (i.e. non-redundant) 150 bp DHSs. FDR 1 % peak calls from all primary tubule and RCC 38 million-tag-subsampled datasets were merged by keeping positions covered by peaks from at least three datasets. Regions where multiple overlapping peaks produced a large contiguous stretch of peak coverage were resolved to multiple, non-overlapping 150-bp segments using a sliding-window approach to find the 150-bp segments of highest coverage within the larger contiguous region.

### Copy-number correction of DNase data

We utilized the “copynumber” package in R to identify genomic regions likely to be subject to copy-number alterations in our RCC samples, with the goal of correcting DNase cleavage counts accordingly so that differences between RCC and TUB samples were more likely to be driven by changes in TF occupancy than by altered copy number. Using the log2-normalized fold-change (RCC/TUB) of DNase tag densities within master list DHSs, we segmented the genomes of all three patient samples (discontinuity parameter gamma = 140). We classified regions whose absolute fold-change were at least twice the median as copy-number variable (Patient 1 = 22 regions, Patient 2 = 26, Patient 3 = 32), and used the mean value of the segment as a scaling factor for raw DNase read counts in those regions for the RCC samples. This analysis detected both 3p loss and 5q gain (confirmed by karyotyping of these patient samples) as well as several focal copy number changes.

### Identification of differential DHSs

We utilized the DESeq2 software package (Love et al. 2014) in R to identify DHSs with significant differences in accessibility between replicate tumor and normal samples, analyzing each patient separately. Copy-number-corrected tag counts meeting a minimum threshold in at least one sample (25) within the master-list DHSs were used as input for DESeq2, and sites that met an FDR threshold of 1% were considered differential DHSs.

### Calling of HIF1/HIF2 binding sites and identification of HIF-occupied DHSs

We used macs2 peaks (FDR 1%) from HIF-1α, -1β, and -2 α ChIP-seq performed in 786-O cells to classify HIF1 and HIF2 binding sites genome-wide. We classified HIF1 binding sites as HIF-1α peaks that overlapped (by at least 50 bp) a HIF-1β peak (1,820 sites) and HIF2 binding sites as HIF-2α peaks that overlapped (by at least 50 bp) a HIF-1β peak (1,243 sites). DHSs in our master list were classified as HIF-positive if they overlapped a HIF1 or HIF2 binding site by at least 37 bp (25% of DHS width).

### Calculation of gene expression changes and GO term enrichment

Gene expression fold-changes were calculated as the log2 ratio of FPKM values for RCC / TUB (0.001 was added to each FPKM value to control for zero values). For each patient, genes with FPKM ≥ 1 in fold-change ≥ 1.5 in RCC were classified as ‘up-regulated’, the converse criteria were used to classify genes as ‘down-regulated’. All other genes were classified as ‘non-changing’, except those with FPKM “ 1 in both TUB and RCC, which were considered ‘non-expressed’. Shared (across all three patients) up- or downregulated gene sets were used (along with the shared non-changing gene list as a background set) as input for the GoRILLA gene ontology enrichment tool.

### Comparisons of regulatory landscapes and differential DHSs among patients

Principal components analysis was performed on log10-transformed DNase I tag densities within master list DHSs (or on FPKM values for RNA-seq data) using the “prcomp” function of R (with center=TRUE and scale=TRUE). Because the master list of DHSs was used to compute differential DHSs for each patient, the DESeq calls (FDR 1%) at each site were used to classify the directionality of change at the same genomic locations across all three patients.

### Connection of HIF binding sites to neighboring differentially expressed genes

We were interested in which genes might be regulated by HIF binding events, and considered clusters of HIF+ DHSs as prime candidates for such connections. To this end, we systematically located clusters of HIF+ DHSs arbitrarily within 12.5 kb of one another, merging neighboring clusters, and examined a 1 Mb region centered on each cluster for genes with altered expression (≥1.5 fold-change) in either our patient samples or TCGA RNA-seq data.

### Survival analyses

Survival analysis based on *POU5F1* expression levels in the legacy TCGA RNA-seq expression data (split evenly into high- and low-expressing groups at the median expression level) was performed using the UCSC Xena web interface (Goldman et al. 2018).

### Uncovering candidate TF drivers of regulatory landscape alterations

Transcription factor motif models were curated from TRANSFAC (version 11) (Matys et al. 2006), JASPAR (Bryne et al. 2008), and a SELEX-derived collection (Jolma et al. 2013). Instances of transcription factor recognition sequences in the human genome were identified by scanning the genome with these motif models using the FIMO tool (Grant et al. 2011) from the MEME Suite version 4.6 (Bailey et al. 2009) with a 5^th^ order Markov model generated from the 36 bp “mappable” genome used as the background model. Instances with a FIMO *P*<10^-4^ were retained and used for subsequent analyses.

To obtain a “family-level” representation of TF recognition sequences, individual motif models used in the genome-wise FIMO scans were compared in a pairwise fashion using the TOMTOM (Gupta et al. 2007) tool from the MEME Suite version 4.6 (Bailey et al. 2009) with the parameters “-dist kullback -query-pseudo 0.1 -target-pseudo 0.1 -text -min-overlap 0 -thresh 1” and the same 5^th^ order Markov model described above as background. Pairwise comparisons were then hierarchically clustered using Pearson correlation as a distance metric and complete linkage. The resulting trees were cut at a height of 0.1 to select clusters of highly similar motifs.

Motif enrichments were calculated by using a custom Python script to count the number of DHSs that contain a “family” motif (i.e. contained an instance of any motif model within a cluster of highly similar motif models). For a given analysis, these counts were compared between a “foreground” set of DHSs (*e.g*. shared DHSs with increased accessibility in RCC) and a “background” set (*e.g*. all other DHSs) and significance was determined using the hypergeometric distribution and subsequent Bonferroni correction of p-values.

Because motif enrichment was computed using family-level representations of TF recognition sequences, we aimed to uncover which member(s) of the POU family might be driving changes in the regulatory landscape of RCC by examining our and TCGA’s RNA-seq data for all members of the POU family with a significant enrichment signal.

### Data access

All primary and uniformly processed sequence data generated in this study are available at the NCBI Gene Expression Omnibus (GEO; http://www.ncbi.nlm.nih.gov/geo/) under accession number GSE117324. We recently performed a separate and non-overlapping analysis of the tubule data sets included in this study in comparison to human kidney glomerular outgrowth cultures and cultured podocytes (*manuscript in revision*). Those data have also been deposited at GEO with accession number GSE115961.

## Author Contributions

SA conceived of the project, procured and processed specimens and designed and performed experiments. KTS, CPM and SA performed experiments and interpreted data. MT performed and interpreted the POU5F1 tissue microarray immunohistochemistry study. SA, KTS, JDV, AR, ER, SJN and EH performed analyses, data interpretation and visualization. RS, AJ and JN processed and curated sequencing data and imported external datasets. DB, MD and DD processed samples for DNase-seq and RNA-seq. RS, MF, MB and RK codified sample metadata and submitted datasets to public repositories. JM and HR-B provided 786-O reagents, interpreted data and edited the manuscript and figures. YZ contributed to experiment design, interpreted data and edited the manuscript. JH supported the study in part, interpreted data and edited the manuscript. SA and KTS primarily wrote the manuscript and all authors edited the manuscript and figures for content and clarity.

## Acknowledgments

SA was supported in part by a Damon Runyon Cancer Research Foundation Fellowship (DRG 114-13). We would like to thank John A. Stamatoyannopoulos whose ENCODE grant from NHGRI (U54HG007010) supported sequencing and data processing for this project. This project was also supported by NCATS grants to JH (5UH3TR000504 and 1UG3TR002158), a NCI Cancer Center Support Grant (P30CA015704) to the Fred Hutchinson Cancer Research Center/University of Washington Cancer Consortium and by an unrestricted gift from the Northwest Kidney Centers to the Kidney Research Institute. We would also like to thank Dr. Kimberly Muczynski and the Kidney Research Institute at the University of Washington for assistance with patient consenting and tissue procurement. We would like to thank Magdalena Skipper and John A. Stamatoyannopoulos for advice on data visualization, figure layout and for editing the manuscript for clarity.

